# Ecological Specialization and Diversification in Birds

**DOI:** 10.1101/2020.06.13.142703

**Authors:** Nicholas M. A. Crouch, Robert E. Ricklefs, Boris Igić

## Abstract

Ecological specialization is widely thought to influence patterns of species richness by affecting rates at which species multiply and perish. Quantifying specialization is challenging, and using only one or a small number of ecological axes could bias estimates of overall specialization. Here, we calculate an index of specialization, based on seven measured traits, and estimate its effect on speciation and extinction rates in a large clade of birds. We find that speciation rate is independent of specialization, suggesting independence of local ecology and the geographic distributions of populations that promote allopatric species formation. Although some analyses suggest that more specialized species have higher extinction rates, leading to negative net diversification, this relationship is not consistently identified across our analyses. Our results suggest that specialization may drive diversification dynamics only on local scales or in specific clades, but is not generally responsible for macroevolutionary disparity in lineage diversification rates.

---

Ecological specialization can be expressed in terms of limits to the range of resources utilized, and environments occupied, by a taxon. Understanding how competition and specialization influence species diversification has been an important goal for ecologists (Forister et al., 2012; Glor, 2010). However, the multidimensional nature of ecological distributions and resource use makes the quantification of specialization challenging. New approaches have benefited from increasing availability of data and more refined quantitative analyses (Devictor et al., 2010; Forister et al., 2012; Poisot et al., 2011). Historically, analyses of specialization have focused on a single aspect, or ecological axis, of species’ ecology. Ecological axes define both the resources exploited by individuals of a species and the environmental conditions under which they live (Devictor et al., 2010; Futuyma and Moreno, 1988; Hutchinson, 1957; Poisot et al., 2011). Focusing on a single axis, such as morphological specialization, could bias interpretation of overall specialization when different ecological axes are important to different species, and multiple axes are important to any given species. This has been appreciated almost since the emergence of ecology as a discipline, when, for example, Grinnell (1917) described the California Thrasher (*Toxostoma redivivum*) as being “*relatively omnivorous*” with respect to diet, but having specific habitat associations reflecting a dependence on cover for protection. More recently, Poisot et al. (2011) noted that a species might “…*simultaneously be a generalist on one axis and a specialist on another*,” confirming that multiple ecological axes must be interpreted concurrently if the objective is an overall measure of specialization. Considering a broad range of ecological axes may therefore help to avoid subjectivity regarding definitions of specialization (Futuyma and Moreno, 1988). This goal has been placed within reach by ever-increasing availability of data on a wide range of species.

Specialization might affect clade richness by influencing the rate at which lineages multiply and become extinct (Bennett and Owens, 2002; Owens et al., 2005; Phillimore et al., 2006; Salisbury et al., 2012; Schluter, 2000). For example, flexible feeding habits might facilitate colonization of new regions more easily, leading to allopatric speciation (Mayr, 1963). Conversely, limited dispersal associated with specialization might also promote allopatric speciation (Salisbury et al., 2012). How such hypotheses are tested has also changed along with analytical advances (reviews by Hernández et al., 2013; Ng and Smith, 2014). The effect of specialization on diversification has been tested through comparison of branch lengths among differently specialized clades (Miller and Crespi, 2003), sister-clade comparisons (reviewed in Tilman, 2008), and Bayesian Mixed Models (Salisbury et al., 2012). If particular characters influenced speciation and/or extinction rates, however, the shape of the tree would not be independent of these characters (Maddison, 2006). Therefore, failing to consider the effect of character states on branching patterns might lead to biased estimates of diversification rate (Maddison, 2006). This issue has stimulated the development of several related methods for testing the effect of character states on speciation and extinction (FitzJohn, 2010; Maddison et al., 2007), although care is required in their application, as model misspecification can profoundly alter results (Goldberg and Igić, 2008; Pennell et al., 2015; Rabosky and Goldberg, 2015).

In this study we ask whether rates of lineage splitting (speciation) and/or extinction in birds are affected by specialization, when defined over multiple axes of species ecology. To address this question, we quantified specialization for 1039 species of land birds in the large and well-sampled monophyletic group Telluraves (Yuri et al., 2013; Jarvis et al., 2014). For each species sampled, we assembled data pertaining to seven ecologically relevant axes, described below, before synthesizing the axis scores into a single metric for specialization. We estimate the relationship of this metric to diversification in a phylogenetic framework, and assess the robustness of our results to specific sources of error.

## Methods

We chose to sample within Telluraves, as this group shows pronounced asymmetry among lineages with respect to ecology and species richness. The Telluraves comprise most land birds with altricial (dependent) development. Species richness ranges from the hyper-diverse passerines (Passeriformes), which account for approximately two-thirds of all birds, to the monotypic order Leptosomiformes (cuckoo-roller). We sampled species from nine orders to cover a range of ecological relationships. These orders : Accipitriformes (hawks and eagles), Bucerotiformes (hornbills and allies), Coliiformes (mousebirds), Coraciiformes (kingfishers, todies, rollers, bee-eaters, and motmots), Falconiformes (falcons), Leptosomiformes (cuckoo-roller), Passeriformes (passerines, or perching birds), Piciformes (woodpeckers and allies), and Trogoniformes (trogons).

We chose ecological axes to provide an overall assessment of specialization, and with a practical constraint of having data available, for a large number of species in each of the taxonomic orders considered. Some ecological axes were more limiting than others. Most notably, clutch size was the most limiting ecological axis, as these data can be extremely hard to obtain owing to remote nesting locations and species nesting in cavities. Each axis can be considered either an ‘Eltonian’ (e.g., morphology) or ‘Grinnellian’ (e.g., habitat) component of specialization (Devictor et al. 2010). Because of these differences, the definition and quantification of specialization varies among axes. The manner of quantifying specialization on each axis was decided *a priori*, with each method having been used in previous studies, with the exception of clutch size specialization. Species were sampled globally but are concentrated in the regions of highest avian diversity (Figure S1).

### Morphological specialization

In this study, we quantified morphological specialization as the position of a species in functional space (Devictor et al., 2010). More ‘generalist’ species are, by definition, more centrally located in the functional space (Belwood et al., 2006; Losos et al., 1994; Mouillot et al., 2007; Ricklefs, 2012). Conversely, species located peripherally are considered more specialized, as these species have morphologies that might constrain movement and feeding in a particular manner, on particular substrates, or on particular dietary items (Figure S2).

We quantified the metric of morphological specialization based on measurements of museum specimens in the collections of the Field Museum of Natural History, Chicago. For all species, we took seven external measurements: wing, tail, tarsus, mid toe, beak width, beak depth, and beak length. These are standard avian measurements of overall size and shape, which have been shown to correspond with ecological difference between taxa (Karr and James, 1975; Miles and Ricklefs, 1984; Miles et al., 1987; Ricklefs and Miles, 1994; Ricklefs and Travis, 1980). For example, the relative lengths of the wings, tail, tarsus, and toes of the White-breasted Nuthatch (*Sitta carolinensis*) demonstrate a particular predisposition for foraging on tree trunks (Miles and Ricklefs, 1984).

We measured one male and one female of each species, with the average of the two being the species value for each trait. The degree of sexual dimorphism within these taxa is dwarfed by interspecies variation, with previous work showing that species’ position in morphological space is unaffected by using sex-specific measurements (Ricklefs, 2012). We log-transformed the measurements to normalize the distribution of the variation (Ricklefs, 2005), and performed a principal components (PC) analysis based on the covariance matrix.

The combination of all PC axes creates a seven-dimensional morphological space, the bounds of which are determined by the morphology of all the species included in the study. The position of a species within the space is determined by its score on all seven PC axes (mean = 0, standard deviation = 1), with morphological specialization defined as the distance between a species location and the center of the morphological space (Belwood et al., 2006; Losos et al., 1994; Mouillot et al., 2007; Ricklefs, 2005, 2012). This is the Euclidean distance, and is calculated as the square root of the sum of the squares of each of the seven normalized PC scores (Ricklefs, 2012).

### Dietary Specialization

We compiled diet information for each species from the Handbook of the Birds of the World (Del Hoyo et al., 2008). Although potentially more accurate approaches exist for quantifying dietary specialization, for example, analysis of gut contents, such data are not available at the taxonomic scale of this study. We documented every reported prey item for each species, even if it was stated as being exploited infrequently. Recording prey regardless of reported frequency minimizes a downward bias in prey diversity among less extensively studied species. We categorized each prey item with respect to class-level taxon prior to calculating dietary specialization. Classifying prey items by taxonomic class, rather than a lower-ranked taxon, generally increases calculated food-item overlap between species (Schluter, 2000), including cases in which the range of exploited items of a more specialized species is included within that of a less specialized species (Futuyma and Moreno, 1988). We created additional categories for classifying dietary items if taxonomic rank was not reported; for example: fruits and seeds were two such categories. Dietary specialization was calculated as the inverse of a standard index based on proportions; Σ(*p*_*i*_)^2^(Devictor et al., 2010; Levins, 1962; Simpson, 1953), where *p*_*i*_ is the proportion of each prey category in the diet. This is a versatile measure of diversity, expressed in terms of the equivalent number of equally common prey items (see MacArthur, 1972), where lower scores represent species with more specialized diets. We calculated a single value for dietary specialization for each species, incorporating any variation that might occur through the year or between populations.

### Clutch-size specialization

The ability of a species to manipulate its clutch size provides an assessment of specificity in a key life history trait. Clutch size can be altered in response to long-term selection, including feedback from climate change (Gienapp et al., 2008), as a density-dependent response (Both et al., 2000) to population pressures or resources, or as a short term response to predation risk (Fontaine and Martin, 2006) or variation in the food supply. Clutch specialization can be quantified by variation in the number of eggs laid per nest (clutch size), expressed as a proportion of the maximum observed clutch size of a species. We calculated clutch specialization for each species (data from Handbook of the Birds of the World) as log(*R*)/log(*M*), where *R* max(eggs) - min(eggs) for the species and *M* is max(eggs) for the species. Clutch specialization ranges from 0 (maximum clutch specialization) to 1. Species with no recorded variation in clutch size, e.g., single-egg layers, received a score of 0.

### Abiotic conditions

We incorporated four abiotic factors into our index of specialization: habitat complexity, temperature range, rainfall range, and altitudinal range within the distribution of each species. We assessed habitat complexity using the Normalized Difference Vegetation Index (NDVI, Hurlbert, 2004; Jetz and Rahbek, 2002; Tucker, 1979). NDVI uses multi-spectral satellite data to monitor the vigor of plant growth, vegetation cover, and biomass production, with higher values corresponding to more complex vegetation cover. We obtained these data from NASA’s Earth Observing System Data and Information System (EOSDIS, 2009). We downloaded temperature, precipitation, and altitude data from the WorldClim database (Hijmans et al., 2005, using the mean layer at 30 second resolution). For each of the abiotic factors, values were extracted from across the range of each species, which, for migratory species, included both the wintering and breeding ranges. We defined specialization as the range of conditions experienced across a species’ range (maximum - minimum values), except for habitat complexity which was defined as the mean NDVI value. The mean NDVI value reflects physical restrictions that habitat structure places on species ecology. Specifically, species that inhabit a more open environment likely experience fewer physical restrictions, for example in moving through the landscape. Conversely, a heterogeneous environment may require a species to perch on different substrates, identify forage among a wider array of resources and substrates, etc.

Species distributions for determining climatic conditions were obtained from BirdLife International and NatureServe (2012). These data provide a broad overview of species distributions, with information on each species gathered from multiple sources including field guides and individual species accounts. Climatic variables were calculated across the extent of species’ ranges. It must be noted that species may only use specific subsets of these total ranges, i.e., have more specific habitat requirements. This may impact the accuracy of NDVI scores in particular; however, more detailed range data are not available at the taxonomic scope of this study. Each of the abiotic variables was converted to raster format in ArcMap (ESRI, 2011) before being analyzed in R (R Core Team, 2013).

### Calculating specialization scores

The specialization score for each species depended on its degree of specialization on each of the seven ecological axes: morphology, precipitation range, altitude range, temperature range, clutch flexibility, diet, and habitat heterogeneity. We weighted each axis equally because we had no prior expectation that particular axes contribute differently to specialization. We scaled each axis by centering the values— subtracting the mean axis score from each species score—and then dividing each axis by its standard deviation. This linear scaling approach can be used for both normally and non-normally distributed data. To produce only positive scores on each axis, we added the absolute value of the minimum observation along each axis. Therefore, the species with the smallest value along an axis received a score of 0, with increasing values representing decreasing specialization. Because scores were normalized by the mean and standard deviation, each axis contributed equally to the specialization index. The overall score a species received was the sum of the scores from each axis; thus, highly specialized species tended to be specialized across all seven of the ecological axes considered here (Dennis et al., 2011).

### Specialization, diversification, and phylogeny

We estimated the effect of specialization on speciation and extinction rates using the Quantitative State Speciation and Extinction model (QuaSSE; FitzJohn, 2010), implemented in the R package ‘diversitree’ (FitzJohn, 2012a). QuaSSE estimates speciation and extinction rates as functions of a continuous variable that evolves by a diffusion process. The functions are defined *a priori*, and can be any that describe the relationship between two variables. For this study, we examine constant, linear, sigmoidal, and modal relationships (FitzJohn, 2010). The initial parameters for the modal function produce a hump-shaped distribution. However, as the parameters describe inflection point variance of the normal distribution, the function can take a variety of shapes that correspond to varying rates (FitzJohn, 2010). We first created 16 evolutionary models, each of which represented a pairwise combination of these four speciation and extinction functions. Each evolutionary model can also include a ‘drift’ parameter, representing an independent directional tendency of a trait over time. In this study, a negative value would indicate an increase in specialization along evolving lineages. Adding a drift parameter created an additional 16 models to be tested.

Analyses of character evolution may be confounded when particular trait values are concentrated within a clade that has undergone recent, rapid diversification. Passerine birds exhibit a significantly higher net diversification rate (0.14 Myr) compared to the rest of the avian tree (0.056 Myr; Jetz et al. 2012), and they account for roughly two-thirds of all bird species. To test for potential effects of asymmetric diversification rates, we examined the models defined above, both those with and without the drift parameter, with the phylogeny partitioned between passerine and non-passerine species. Partitioning the tree allowed us to estimate independent rates between partitions. The combination of rate functions, with and without the drift parameter, and with and without the tree partition, gave a total of 64 models to be tested (Table 1).

**Table 1:** AIC results of QuaSSE model fitting. Values are AIC followed by AIC*w* scores. AIC*w* were calculated from the full set of 64 models. Speciation and extinction refer to the function used to model each respectively. Drift is a directional term, indicating whether there is a tendency towards an increase or decrease in the character state through time. In partitioned models the phylogeny was split between passerine and non-passerine species, allowing independent parameter estimation for the two groups. The best fitting model is highlighted in bold.

|  |  | Unpartitioned phylogeny |  | Partitioned phylogeny |  |
| --- | --- | --- | --- | --- | --- |
| Speciation | Extinction | No Drift | With Drift | No Drift | With Drift |
| Constant | Constant | 13949.94/0.0 | 13945.43/0.0 | 14277.32/0.0 | 14282.68/0.0 |
| Constant | Linear | 13949.72/0.0 | 13945.86/0.0 | 14279.64/0.0 | 14140.43/0.0 |
| Constant | Sigmoid | 18820.23/0.0 | 14340.47/0.0 | 17177.59/0.0 | 17098.41/0.0 |
| Constant | Modal | 16620.34/0.0 | **13907.41/0.9** | 15601.31/0.0 | 15538.01/0.0 |
| Linear | Constant | 13956.64/0.0 | 13947.99/0.0 | 14292.63/0.0 | 14305.70/0.0 |
| Linear | Linear | 13957.95/0.0 | 13952.83/0.0 | 14264.36/0.0 | 14257.82/0.0 |
| Linear | Sigmoid | 19247.64/0.0 | 18786.25/0.0 | 19602.44/0.0 | 19580.39/0.0 |
| Linear | Modal | 13952.22/0.0 | 13936.03/0.0 | 14203.56/0.0 | 14181.68/0.0 |
| Sigmoid | Constant | 14185.77/0.0 | 14079.18/0.0 | 14561.98/0.0 | 14516.03/0.0 |
| Sigmoid | Linear | 14187.77/0.0 | 14110.31/0.0 | 14581.23/0.0 | 14530.43/0.0 |
| Sigmoid | Sigmoid | 19660.87/0.0 | 16709.25/0.0 | 19761.23/0.0 | 18752.58/0.0 |
| Sigmoid | Modal | 14187.02/0.0 | 14029.08/0.0 | 14389.20/0.0 | 14422.49/0.0 |
| Modal | Constant | 13972.28/0.0 | 13950.08/0.0 | 14153.11/0.0 | 14298.39/0.0 |
| Modal | Linear | 14074.37/0.0 | 13956.54/0.0 | 14405.29/0.0 | 14408.23/0.0 |
| Modal | Sigmoid | 19828.85/0.0 | 14154.98/0.0 | 20275.02/0.0 | 20025.39/0.0 |
| Modal | Modal | 14061.35/0.0 | 13947.24/0.0 | 14243.76/0.0 | 14241.28/0.0 |

Phylogenetic data for model fitting and parameter estimation were sampled from Jetz et al. (2012). To include all extant species of birds, those authors created a species-level phylogeny with sequence data. Species lacking sequence data were placed by Jetz et al. (2012) on this phylogeny using species taxonomy. The branch lengths for these species were drawn randomly from a pure birth process, resulting in variation in both topology and branch lengths for species both with and without genetic data. Accordingly, parameter estimation of speciation and extinction rates was calculated across ten randomly selected phylogenies. To estimate whether the variation in tree structure in the ten sampled phylogenies encompassed variation across a wider set of phylogenies, we calculated pairwise distances between our phylogenies using three metrics (Colijn and Plazzotta, 2018; Kuhner and Felsenstein, 1994; Robinson and Foulds, 1981), calculated using the R package *phangorn* (Schliep et al., 2017). In each case, we compared the variation among our ten phylogenies against the pairwise distances of 100 phylogenies sampled from Jetz et al. (2012).

Model fitting was performed by calculating the likelihoods of the estimated parameters of each model describing the process that shaped both the branching times in the phylogeny, and the distribution of the trait at the tips. Akaike Information Criterion (AIC) scores (Akaike, 1974) were generated for each model, penalizing the likelihood score by the number of parameters in the model. Comparison of AIC scores shows the relative support of each model, and can be used to calculate the probability (Akaike weights) of any given model being the best of those tested. Here, we calculated the Akaike weight (AICw) of each model from the combined AIC scores of all 64 models using the R package *qpcR* (Spiess, 2018).

Estimating character trait-dependent diversification rates for binary traits can produce false positive results (Rabosky and Goldberg, 2015). To estimate the propensity for our continuous character to give false positive results, we permuted the calculated species specialization scores 75 times to generate simulated random data sets. Using these data, and a single sampled phylogeny, we carried out model fitting for a single state-dependent model, and for a single state-independent model, using QuaSSE. A state-dependent model allows the speciation and extinction rates to vary depending on the trait value. Here, the state-dependent model specified a linear function for both the speciation and extinction rates. Conversely, in a state-independent model, neither the speciation nor the extinction rate changes with changes in the trait value (i.e., both take a constant value); such a model should be preferred for a neutral character. Using AIC scores to compare the fit of the two models, with the error rate defined as the frequency with which a state-dependent model was spuriously estimated to explain the data better than a state-independent model. This rate reflects a degree of inference error in these analyses and, although it is an aspect of type I error, it is not the only possible source of false positive results (Rabosky and Goldberg, 2015).

### Assessing sensitivity to definitions of specialization

We also performed a number of additional analyses to determine how sensitive our results are to different treatments of the data. First, we analyzed each of the seven ecological axes independently to investigate whether the use of a compound metric of specialization is potentially obscuring the signal of one or several individual traits affecting speciation and/or extinction rates. Next, we derived different compound metrics that differed either in the number of ecological axes included, or in treatment of the data. The first of these alternative compound metrics excluded the three climate axes (range in temperature, altitude, and precipitation) as these may co-vary with range size and so may bias the results. Next, we derived a metric that excluded the morphology axis as, in the principal analysis, we did not perform a a phylogenetic PCA. Finally, we calculated a metric that included all seven ecological axes, but with the morphological specialization scores calculated using phylogenetic PCA (using the R package *phytools*, Revell, 2012) After morphological specialization scores were calculated using the phylogenetic PCA data, they were combined with the remaining six axes as described above.

In all of these sensitivity analyses, based on the results of the main analysis, we fitted three models to analyses of individual ecological axes, namely: (i) where both speciation rate and extinction rate were independent of the character value (ii) where both speciation and extinction took a variable rate, and (iii) where the speciation rate was held constant while the extinction rate was allowed to vary. Additionally, all of the model sensitivity analyses were performed on a maximum clade credibility (MCC) phylogeny created from the posterior distribution of Jetz et al. (2012).

### Alternative phylogenetic analyses

We supplemented the QuaSSE analyses by comparing the individual trait axis values and the three compound metrics to speciation and extinction rates estimated using BAMM (Bayesian analysis of macroevolutionary mixtures, Rabosky 2013, 2014; Rabosky et al. 2014). We used BAMM both as an alternative analytical technique, and also to accommodate for potential issues caused by incomplete sampling. Specifically, we used speciation rate estimates derived from Harvey et al. (2017) who analyzed a maximum clade credibility tree from the genetic-only analysis of Jetz et al. (2012). This analysis included 6,670 species, incorporating 67% sampling rate in BAMM, thus ensuring highly robust speciation rate estimates. We tested for correlations with the trait values using structured rate permutations of phylogenies (STRAPP; Rabosky and Huang, 2016), which samples speciation rate estimates from the posterior distribution of the BAMM analysis and tests for correlations with trait values. We sampled 1000 samples from the posterior distribution, log-transforming the values prior to analysis. In total we performed 20 STRAPP analyses.

### Independent assessment of extinction risk

Some potential issues arise from estimating extinction rates from phylogenetic data (Rabosky, 2010). Therefore, as an independent approach to estimating extinction risk, we compared our metric of specialization against International Union for the Conservation of Nature (IUCN) extinction risk categories. The IUCN codes cover a range of possible extinction risks, including anthropogenic influences. Risk assignments also incorporate the range size of species, meaning that the climatic variables in our metric of specialization might bias any correlation with IUCN scores. Therefore, we tested for a relationship between IUCN scores and the four-axis definition of specialization that excludes the climatic variables. Finally, we performed the same comparison using the definition of specialization that incorporates phylogenetically-corrected morphological specialization scores.

## Results

We collected data for 1039 species, representing 93 families of birds (Table S1, S2). Our metric of specialization for these species ranged between 1 (White-throated Grasswren, *Amytornis woodwardi*), the most specialized species in this study) and 25.2 (Red-backed Shrike, *Lanius collurio*) with a median value of 10.3 and mean of 10.2. Figure 1 illustrates the contributions of individual specialization proxies to the overall measure of specialization for a selection of species. Phylogenetic signal for the trait was generally low, with a range of values throughout the tree (Figure S3).

**Figure 1:**
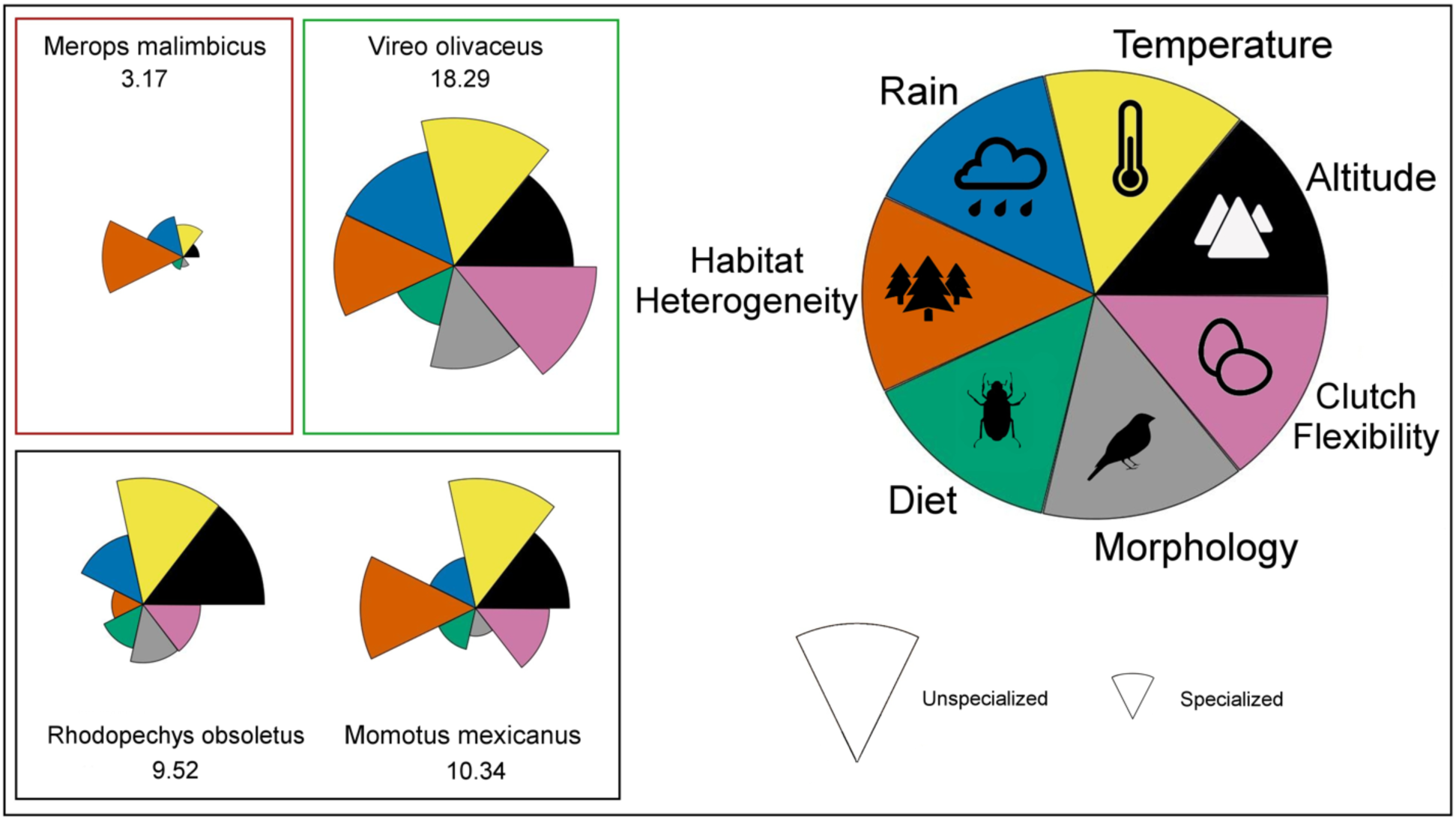
Examples of how calculating specialization as the sum of the score on each ecological axis, represented by different colored pie sections, generates variation in the overall impression of specialization for a species. The size of each section corresponds to how specialized a species is on that ecological axis, with a smaller size representing greater specialization. The numbers below each species correspond to how specialized they are estimated to be based on all seven ecological axes. Shown here is one species estimated to be highly specialized (Rosy bee-eater, *Merops malimbicus*), one unspecialized species (Red-eyed vireo, *Vireo olivaceus*) and two intermediary species (Desert finch, *Rhodopechys obsoletus* & Russet-crowned motmot, *Momotus mexicanus*).

### Specialization and diversification

The group of best-fitting QuaSSE models included a “drift” term, representing a general trend over time, and no difference (partition) between passerine (songbirds) and nonpasserine birds (Table 1). The best model included a single speciation rate, and a varying extinction rate (Table S3, ΔAIC = 28.62 to second best-fitting model). The inferred extinction rate increases with greater specialization, leading to negative net diversification (Figure 2). This model also incorporates a negative drift parameter, pointing to a general increase in specialization over time. This result does not appear to have been influenced by our sampling (supplementary material, Figure S4).

**Figure 2:**
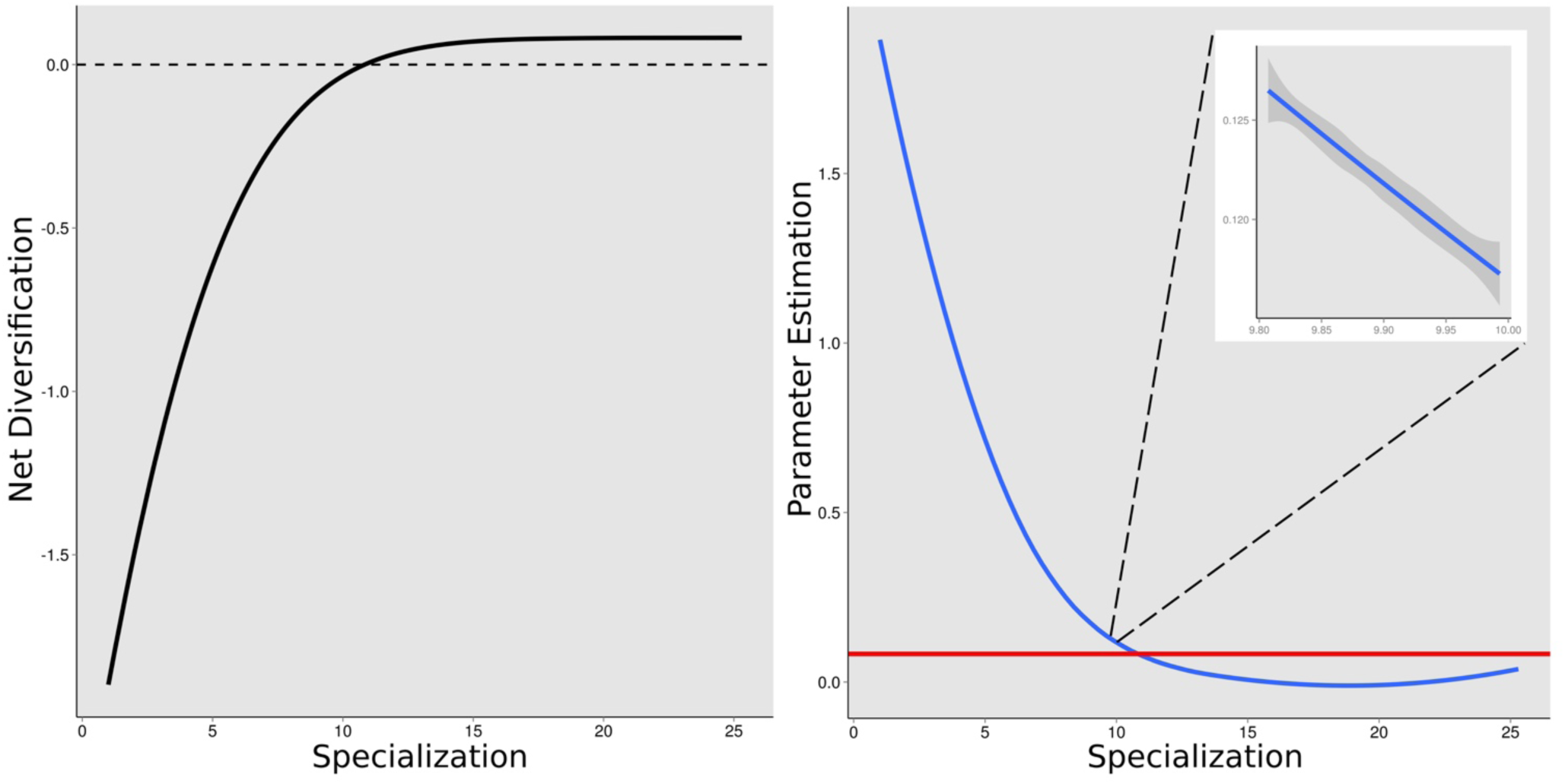
Relationship between net diversification rates and our index of specialization. (Left panel) More specialized species, i.e. those with smaller scores, are estimated to have a negative net diversification rate. (Right panel) Net diversification broken down into speciation (red) and extinction (blue) rates. The confidence limits for speciation and extinction rates are contained within the width of the respective lines. The inserted image shows a magnified section of the extinction rate function to illustrate the variance between the ten sampled phylogenies.

Topological variation in the sampled phylogenetic data had a minimal impact on parameter estimation. Variation would be accentuated if the phylogenetic data contained a large number of species that lacked genetic data in the analysis of Jetz et al. (2012), but few of the sampled species lacked genetic information (148 species, 14% of tips), and those species lacking genetic data were placed non-randomly using constraint parameters by Jetz et al. (2012). The disparity in the sampled phylogenies is, however, also representative of variation seen in a wider set of phylogenies from Jetz et al. (2012), with only a more extreme region of tree space not incorporated (Figure S5). There was low variance in estimated parameter values among phylogenies (Table S4), and very low variance in the resulting shape of the rate functions (Figure 2).

None of the 20 STRAPP analyses using speciation and extinction rates estimated using BAMM produced significant correlations (Figures S6 and S7). For the analyses of individual trait axes there was a slight tendency for positive relationships with both speciation and extinction rates, but none were significant at the 0.05 level. One variable (altitude), had a *p*-value of 0.05, but a very low correlation estimate of 0.15. Correlations with the compound trait metrics were all extremely low, with the highest correlation with speciation rates being 0.15.

### Comparison of specialization and IUCN scores

We recovered different patterns when comparing different metrics of specialization with IUCN risk categories (Figure 3). When we compared our principal metric of specialization we found the expected trend that more at-risk species, those listed as critically endangered (CR) or endangered (EN), were estimated to be significantly more specialized than taxa in the other categories. When the compound metric incorporating a phylogenetic-PCA definition of morphological specialization was tested, a similar pattern was recovered. However, when we tested a metric combining four ecological axes, excluding those variables that might be correlated with range size, we found little difference between the five IUCN categories.

**Figure 3:**
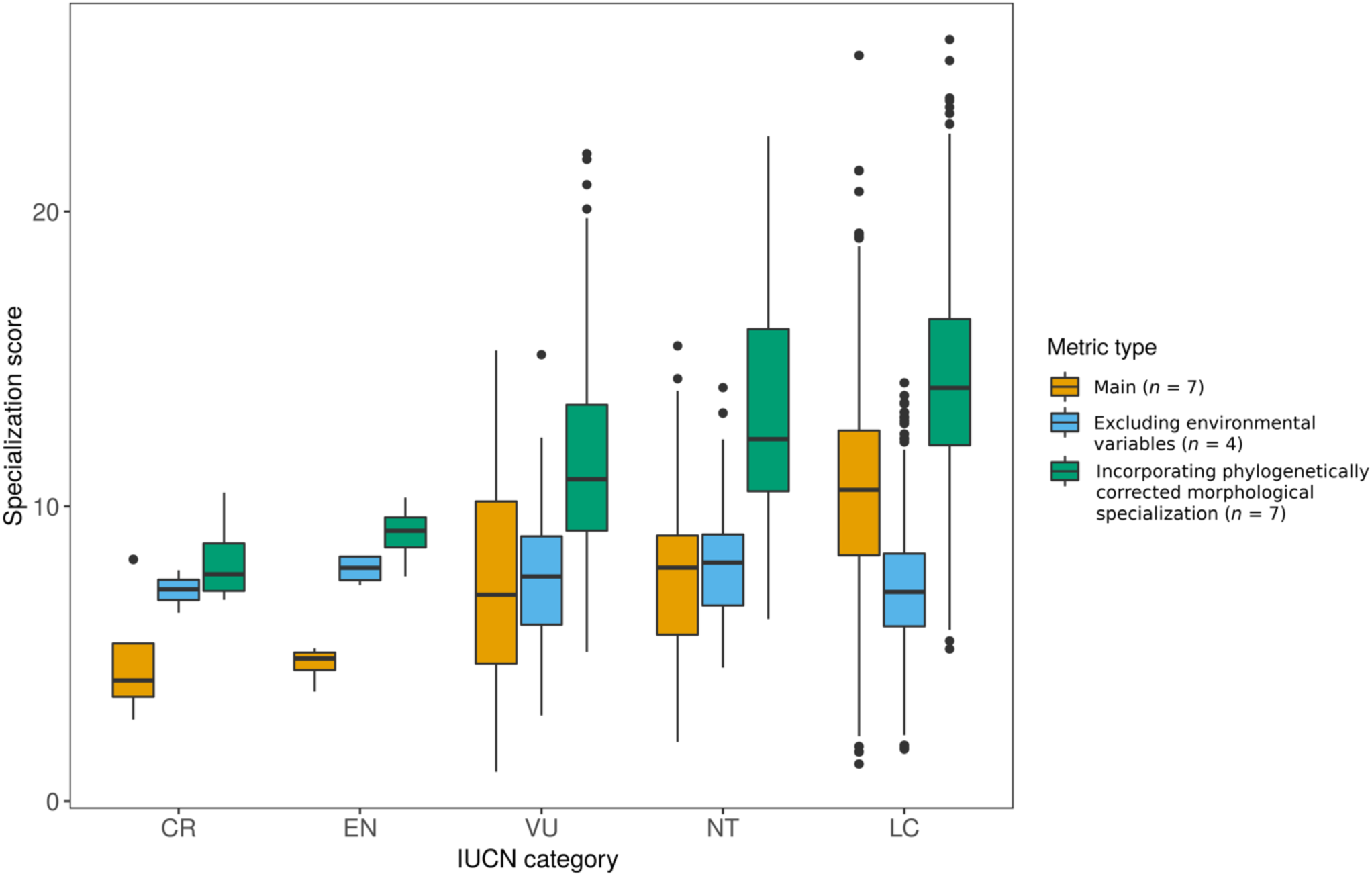
Specialization score vs IUCN Red List categories, corresponding to the estimated extinction risk for species. (A smaller specialization score corresponds to a more specialized species.) IUCN data were available for 1038 species. Categories are ordered by extinction risk: CR = critically endangered (*n* = 3), EN = endangered (*n* = 4), VU = vulnerable (*n* = 27), NT = near threatened (*n* = 43), LC = least concern (*n* = 961). Colors represent different metrics used for defining specialization, with n referring to the number of ecological axes in each definition.

### Sensitivity analysis results

The rate of recovering spurious associations between a neutral character and changes in speciation and/or extinction rate appear to be modest. Only 7 instances (9% of simulations) erroneously favored a state-dependent model over a state-independent model, i.e., when the tip values were randomly re-sampled. This is dramatically lower than the error rate of 77% in a sample empirical analysis of Cetacean (whale) data, in which a continuous size trait was arbitrarily divided into “small” and “large” for a BiSSE analysis (Rabosky and Goldberg, 2015). Machac (2014) suggested that the type I error rate for QuaSSE analyses increases with increasing deviation from constant diversification, particularly when diversification slows over time. Under the greatest deviation from constant diversification (γ = −5), Machac (2014) estimated the type I error for QuaSSE to be 20%. The estimated slowdown in diversification in the phylogenetic data used here was even greater (mean γ = −6.44, 0.03 standard error). Therefore, our data appeared to be less prone to the source of error found by Rabosky and Goldberg (2015) and Machac (2014).

## Discussion

We found that speciation rate in a large sample of avian taxa is independent of a multidimensional measure of specialization. We also found that highly specialized species exhibit elevated extinction rates, leading to negative net diversification rates. Our results seem robust against both phylogenetic uncertainty and spurious correlations between changes in speciation and extinction rates with randomized trait values.

### Defining specialization as a compound metric

In this study we used a compound metric of specialization, as we argue that different ecological axes must be analyzed simultaneously if the overall aim of a study is to equate variation in species-level specialization with variation in speciation and extinction rates. This is because the different aspects of species’ ecology do not act independently – the propensity for a species to split at a given point in time is not dependent on the effect of one ecological axis while ignoring the effects of others. A potential consequence of this approach is that, if some axes increase, and others decrease the diversification rate, these effects may offset each other and result in the rate appearing to be independent of the metric. This seems likely given that many of the ecological axes included here have been suggested to affect speciation rate. However, in our analysis, speciation and extinction rates were estimated to be independent of each individual ecological axis (Table S5). While this further suggests that ecological differentiation and reproductive isolation are unrelated, it also highlights the utility of compound definitions. The only association between extinction rate and specialization, consistent with a strong prior expectation, involved the analysis of compound metrics. Nevertheless, variation in results obtained between analyses requires further evaluation.

### Specialization and speciation

We are unaware of many cases in which speciation rate is independent of specialization. Salisbury et al. (2012) found that the positive association between net diversification (speciation - extinction) and specialization in their analyses of 739 species of Amazonian birds was diminished when their several metrics of specialization were combined. Our own result might be influenced by the fact that diversity dependence is not accounted for in the models of character evolution (FitzJohn, 2010), with the filling of ecological space likely important to the evolution of specialization. Early in radiations, ecological space is filled rapidly as species exploit novel resources (Schluter, 2000). The rate at which the space is filled depends on the amount of resources available as well as the species present – presumably generalists fill ecological space faster than specialists. Regardless, increasing numbers of species within an environment requires that species partition resources more finely (Hutchinson, 1961), leading to increased specialization. Eventually, as available ecological space becomes full, diversification presumably begins to slow (Phillimore and Price, 2008; Schluter, 2000; Simpson, 1953). Therefore, estimating the effect of specialization on speciation rate could be confounded if specialization were influenced by species diversity. Our sampling of taxa representing highly diverse ecological relationships might have mitigated this problem, as species likely occupy disjunct regions of ecological space. Furthermore, an increasing body of literature suggests that ecological differentiation – presumably associated with increased specialization – is not required for diversification (Mendelson et al., 2014; Nosil and Flaxman, 2010). Therefore, although individual cases emphasize a role for ecological differentiation (i.e., specialization) in species formation (Schluter, 2001, Table 1), our results further suggest that specialization might not be as closely tied to diversification as previously thought (Svensson, 2012).

An historical emphasis, prior to extensive use of phylogenetic comparative methods, relating speciation to ecological differentiation might have resulted from inferring the causes of speciation from biological patterns. For example, more specialized species often exhibit smaller range sizes (Brown, 1984; Holt, 2003; MacArthur, 1972; Williams et al., 2009). This observation culminated in hypotheses linking specialization to speciation rates. For example, reduced ability of species to disperse between habitats increases the isolation and subsequent divergence of allopatric populations (Bryson et al., 2013; Salisbury et al., 2012; Von Rintelen et al., 2010). Reducing species tolerances to fluctuations in abiotic and biotic conditions might also restrict distributions and facilitate diversification (Sheldon, 1993). In such cases, however, no direct link between ecological differentiation and reproductive isolation is established, and doing so is problematic (Price, 2008). Although our data suggest a negative relationship between specialization and range size (Figure S8A), this pattern need not lead to species formation.

Our result may also be biased by a predominance of environmental variables (*n* = 4). As a result, the overall definition of specialization is biased towards Grinnelian aspects of specialization. Additionally, defining specialization as we have done allows individual variables to differentially contribute to the overall definition of specialization. We suggest that additional analyses should evaluate how different assumptions about the choice of ecological axes, and how differential weighting of those axes, affect both our understanding of how specialization is defined, and also its inferred effect on macroevolutionary dynamics. Such efforts will likely be aided by the increasing availability of high-quality life-history data on a diverse range of taxa.

### Specialization and extinction

The estimated relationship between specialization and extinction varied between analyses. In our main analysis the shape of the function relating extinction rate to specialization indicates a high extinction rate for the most specialized species, i.e., those with the lowest specialization scores (Figure 2). Several authors have suggested that specialization should increase extinction rates (e.g., McKinney, 1997) owing, for example, to reduced ability to cope with climatic variation, competition with non-native species, and decreased resistance to novel pathogens (reviewed by Dennis et al., 2011). Although our analysis supports the relationship, extinction rates inferred from phylogenies should not be interpreted as more than general trends (FitzJohn, 2010, 2012b; Rabosky, 2010; Ricklefs, 2007; Stadler, 2012). Moreover, we did not consistently identify a relationship between specialization and extinction, most importantly when the phylogenetic PCA definition of morphology was incorporated into the seven-axis definition (Table S5). Therefore, although any given analysis may confidently recover a relationship, such a result is not necessarily uniformly supported.

Phylogenetic methods for estimating the effect of a trait value on extinction rates can be supplemented by alternative approaches. Here, we compared our different estimations of specialization against IUCN-assigned categories of extinction risk. A comparison of the two metrics of specialization that incorporate seven ecological axes both suggest a relationship with extinction risk; however, the climatic variables incorporated in those definitions are correlated with range size, causing these metrics of specialization to be correlated also (Figure S8A). Removing these variables produces a range-size-independent metric (Figure S8B), which is not identified as being correlated with IUCN categories (Figure 3). Therefore, we can only draw tentative conclusions between specialization (as presented here) and extinction, despite the strong prior assumption of such a relationship.

### Temporal trends in ecological specificity

The estimated “drift” term from the best fitting QuaSSE model from the main analysis was negative, consistent with species becoming increasingly specialized over time. Increasing specialization through time is suggested by theory, as well as being demonstrated in empirical studies (Darwin, 1859; Futuyma and Moreno, 1988; Kelley and Farrell, 1998; MacArthur and Wilson, 1967; MacArthur and Levins, 1967; MacArthur and Pianka, 1966; Nosil, 2002; Nosil and Mooers, 2005; Simpson, 1953). Species become specialized through adaptation to a narrow range of resources (Poisot et al., 2011). This process can be reinforced as increasing specialization leads to reduced morphological and genetic variation (Futuyma and Moreno, 1988; Poisot et al., 2011), which can encumber expansion of a range of resources (e.g., Paull et al., 2012, and references therein).

Increasingly, however, examples surface of taxa that do not follow this ‘generalist-to-specialist’ pattern (Colles et al., 2009). Because specialization appears to be labile, one might ask whether its estimated directional trend is an artifact of a particular analytical approach. Specialization might decrease following expansion of ecological opportunity resulting from, for example, island colonization, removal of competing species, and the acquisition of novel trait(s) (Glor, 2010; Losos, 2010). Such instances of decreased specialization might not be picked up in phylogenetic analyses of character evolution at the taxonomic scale of this study.

Trends in trait values are also likely affected by the analytical approach taken. Here, analysis of a compound metric of specialization, where a phylogenetic PCA is performed prior to the calculation of morphological specialization scores, results in the ‘drift’ parameter estimated to be positive, indicating a decrease in specialization through time. This is likely driven by the fact that passerines, one of the most derived groups within Telluraves, are concentrated near the center of morphospace. Therefore, a lack of a phylogenetic correction will result in estimated directional evolution of species’ trait values. Although the estimated ‘drift’ values were small in the corrected metric, approximately half that of the rate of evolution, this result highlights how different treatments of the data can result in significant differences in results. Given the variance in results obtained here we are hesitant to draw strong conclusions, and suggest that this will be a fruitful avenue of future research.

## Conclusion

Achieving a comprehensive theoretical, as well as practical, understanding of specialization is difficult considering the multidimensional nature of ecological relationships including both physical and biological aspects of the environment. Our results, based on specialization estimated over multiple ecological axes, present little support for specialization either affecting or following upon macroevolutionary dynamics. Some analyses support specialization affecting the extinction rate of lineages, an intuitively appealing generalization, but perhaps obscured by limitations of the data and available methods of analysis. As the estimated speciation rate appears to be independent of our specialization metric, specialization would appear to be a consequence, rather than a cause, of species formation.

